# Occipital tACS does not modulate manual performance but fixation variability under visuo-proprioceptive conflict

**DOI:** 10.64898/2025.12.08.693038

**Authors:** Josephine Gräfe, Peng Wang, Daria Antonenko, Agnes Flöel, Jakub Limanowski

## Abstract

The modulation of oscillations at alpha and beta frequency ranges recorded over sensory cortices has been linked to the top-down weighting of sensory information relative to (competing) information from other modalities. Here, using a virtual reality-based hand-target matching task under visuo-proprioceptive conflict, we tested whether alpha-/beta-tACS over the occipital cortex would improve action performance when conflicting visual movement feedback needed to be ignored in favor of proprioception. Participants had to match a target rhythm with either their unseen real hand (RH) grasping movements, or with the movements of a virtual hand (VH) that displayed their actual movements after a constant time delay. Visual movement feedback was, consequently, either task-relevant (VH task) or a distractor (RH task). In a cross-over, double-blind, within-subject design, we applied either low-intensity (2 mA peak-to-peak) transcranial alternating current stimulation (tACS) at 10 Hz (“alpha”) or 20 Hz (“beta”), or sham, over the occipital cortex during this task. Neither alpha- nor beta-tACS significantly affected hand-target matching performance, but fixation variability in the RH condition significantly decreased during beta-tACS relative to sham stimulation. Thus, despite not directly modulating manual performance, occipital beta-tACS could have facilitated ignoring the mismatching (distracting) visual movement feedback when needed.

## Introduction

The attenuation of low-frequency oscillations recorded over sensory brain regions has been linked to endogenous attention: For instance, when participants attend to visual, auditory, or somatosensory stimulus attributes, a reduced power of oscillations in the “alpha” frequency can be observed at electrodes covering the respective primary or secondary sensory cortices (e.g., Arnal et al., 2011; Foxe et al., 1998; Fu et al., 2001; Foxe & Simpson, 2005; Kelly et al., 2006; Buschmann & Miller, 2007; Schroeder & Lakatos, 2009; Haegens et al., 2012; Gundlach et al., 2024). Similar effects have been reported for oscillations in the low “beta” range (Bauer et al., 2006, 2012; Limanowski et al., 2020; cf. Lee et al., 2013). This alpha/beta suppression in sensory cortices has been interpreted as indicating top-down processes related to the up-weighting of attended sensory information relative to other signals, including competing sensory information from other modalities (Fries, 2001; Arnal & Giraud, 2012; Bauer et al., 2014: Bastos et al., 2015; Donner & Siegel, 2011; Engel & Fries, 2010; Miller & Buschmann, 2013; Spitzer & Haegens, 2017).

This interpretation has also received support from brain stimulation studies using tACS at alpha or beta frequencies. TACS is the application of oscillatory currents on the human scalp, based on the assumption that this modulates ongoing rhythmic brain activity (Kanai et al., 2008; Zaehle et al., 2010; but cf. Rjosk et al,. 2016; Roshchupkina et al., 2020; Lafleur et al., 2021). In particular, previous work suggests that alpha-tACS over occipital or parietal cortices may modulate visual attention (Helfrich et al., 2014; Clayton et al., 2019; Schuhmann et al., 2019; Nakazono et al., 2020; Kasten et al., 2020; He et al., 2022; Hilla et al., 2023; Bergmann et al., 2025). TACS in the beta range has primarily been applied over primary motor cortices in sensorimotor tasks (e.g., Pogosyan et al., 2009; Schutter & Hortensius, 2011; Nakazono et al., 2016). However, beta-tACS has also been found to modulate visual attention similar to alpha; e.g., beta-tACS over the parietal cortex improved performance in a visual crowding task (Battaglini et al., 2020).

We have recently observed that low-beta oscillatory power over posterior electromagnetic sensors was modulated during a sustained attention, hand-target matching task under visuo-proprioceptive conflict (Limanowski et al., 2020): Participants had to match a target rhythm with a right-hand grasping rhythm, with their real hand (RH) hidden from view and a virtual hand (VH) providing visual movement feedback with a constant delay. The target rhythm had to be matched with either the unseen real hand grasping movements (RH task), or the delayed movements of the virtual hand (VH task). Due to the visuo-proprioceptive conflict introduced by the constant delay, only one of the hands could be aligned with the target at a time, while the respective other hand effectively constituted a cross-modal distractor. Thus, the VH task required enhanced visual attention to prioritize visual over (conflicting) proprioceptive movement feedback for hand-target matching (cf. Lajoie et al., 1992; Balslev et al., 2004; Bernier et al., 2009; Heuer & Rapp, 2012; Limanowski, 2022; Wang & Limanowski, 2023). Conversely, the RH task required ignoring visual feedback for proprioceptive hand-target matching. In a simulation study, we demonstrated that a failure to ignore conflicting visual feedback resulted in poorer RH task performance in this kind of task (Limanowski & Friston, 2020b); in line with previous reports that observing movements incongruent with one’s own biases movement execution (Brass et al., 2001; Kilner et al., 2003; Heyes, 2011). Crucially, we observed stronger posterior/occipital beta power during the RH than the VH task; i.e., those low-beta oscillations were suppressed when participants had to use delayed visual movement feedback to guide action, but augmented when they had to ignore vision and guide action by proprioception. This aligned with the aforementioned suggested inverse relation of sensory alpha/beta frequencies with the top-down (i.e., task-dependent) attentional weighting of sensory information.

In the present study, we aimed to test whether modulating occipital alpha or beta oscillations via tACS would, correspondingly, affect behavioral performance in the same visuo-proprioceptive conflict task. Our key hypothesis was that alpha- and/or beta-tACS, relative to sham stimulation, would improve performance in conditions where visual feedback needed to be ignored (i.e., in the RH task), and significantly more strongly so than in conditions where visual feedback needed to be attended to (VH task). In other words, we expected an interaction effect between stimulation type (alpha-/beta-tACS vs sham) and instructed hand modality (VH vs RH task) on behavioral performance. We tested this hypothesis by analyzing the mean and the variability of hand-target matching and fixation, respectively. I.e., we expected both manual performance and the ability to maintain central fixation (as instructed) would indicate the difficulty of performing the task under visuo-proprioceptive conflict, and any related effects of brain stimulation.

## Methods

### Participants

24 healthy, right-handed, non-smoking participants with normal or corrected-to-normal vision (19 female, mean ± SD age: 21.9 ± 2.3 years) completed the study. Participants received 10€/hour or student credit as compensation. The sample size of 24 was chosen based on prior work using a similar task (Limanowski et al., 2020; Limanowski & Friston, 2020a,b). The experiment was approved by the local ethics committee of the University Medicine of Greifswald.

### Task Design

The participants wore a data glove (5DT Data Glove 5 Ultra, https://5dt.com/5dt-data-glove-ultra/, 1 sensor per finger, 60 Hz sampling rate) on their right hand, which was positioned on a small pillow in their lap and occluded from view via a black barber gown. The averaged flexion values of the four glove sensors were fed to a photorealistic virtual right-hand model, presented on a computer screen in front of the participant (see Fig. 1A). We added a constant time delay of 500 ms to the transmitted data; i.e., the VH movements lagged behind the actually executed movements. Thus, we introduced a permanent conflict between visual and proprioceptive hand movement information (i.e., the seen from the felt hand position).

**Figure 1.**
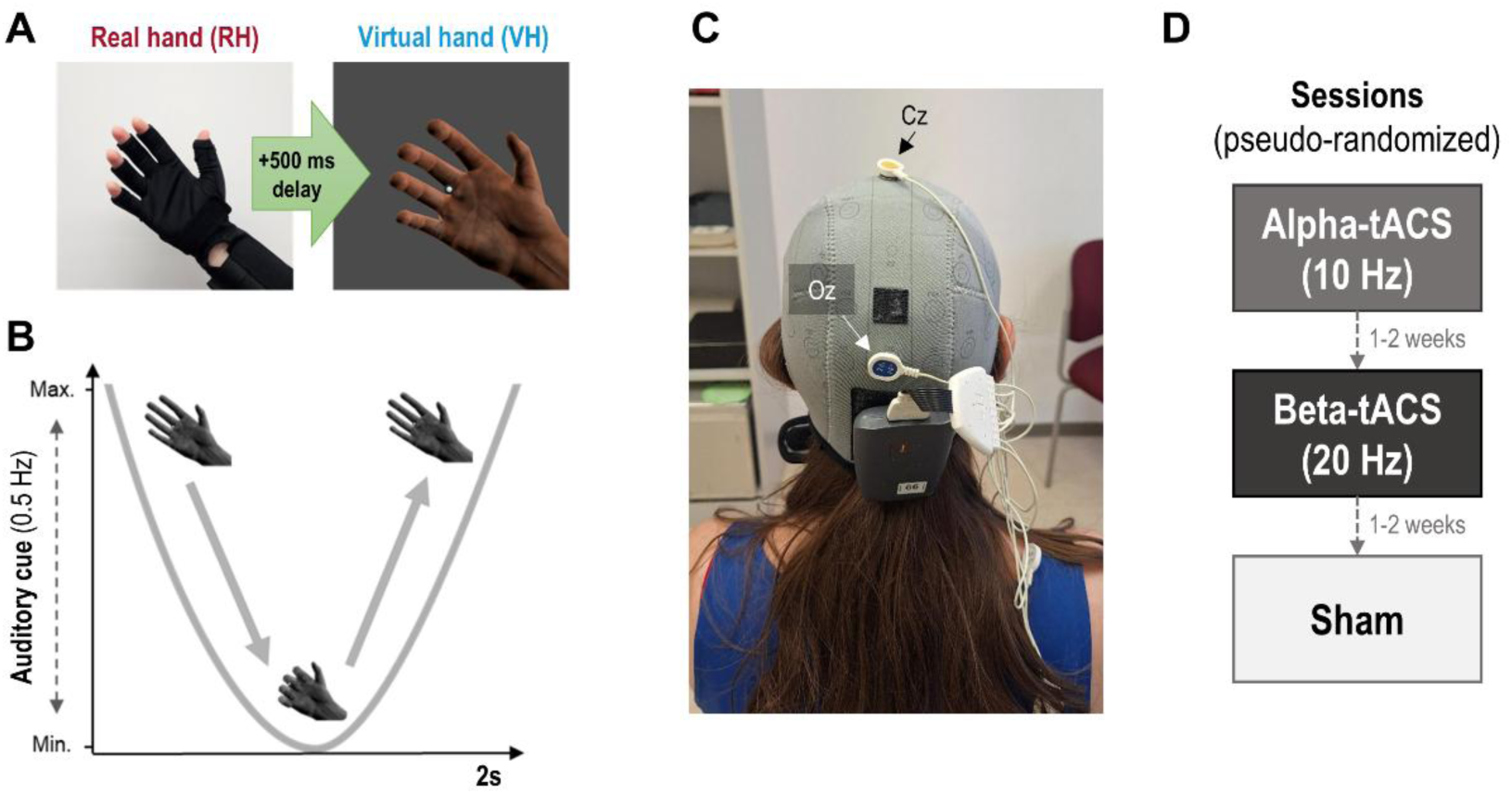
Experimental task and design. A: Participants controlled a virtual hand (VH) model via a data glove worn on their right real hand (RH), which was occluded from view. A constant delay of 500ms was added to the glove data so that the VH was lagging with respect to the participants’ actual hand movements. This introduced a constant visuo-proprioceptive conflict. **B:** The participants’ task was to align recurrent grasping (closing-and-opening) movements with an 0.5 Hz auditory target oscillation (a tone that decreased and increased in volume following an 0.5 Hz sine wave). Thereby participants were instructed to align either the VH or the unseen RH with the target oscillation. Note that, due to the constant delay between the VH and RH movements, only one of the hands could be aligned with the target, while the other one would consequently move out of phase. Thus, the VH task required shifting the RH movements to adapt to lagging visual feedback, while the RH task required ignoring the incongruent (lagging) visual feedback to keep the RH aligned with the target. The hand-target matching task was performed continuously in blocks of 130 s duration. **C:** During the task, we targeted visual cortical oscillations by applying tACS over the occipital cortex (electrode positions: Oz and Cz). **D:** Each participant completed 3 stimulation sessions à 20 min duration, each containing 4 VH and 4 RH task blocks of 130 s each, interspersed with 10 s pauses, in randomized order. The stimulation sessions were completed with 1-2 weeks separation in pseudo-randomized order. Both participant and experimenter were blind with respect to the stimulation conditions.

Participants had to perform continuous grasping (closing-and-opening) hand movements following an auditory pacing signal: a 250 Hz tone changing its volume following a 0.5 Hz sine wave (i.e., one full cycle every 2 s). Thus, when the tone was at its loudest, participants were instructed to have the hand completely open, and when it was silent, completely closed (see Figure 1B). Crucially, participants had to perform this hand-target phase matching task with one of two goals in mind: matching the target’s phase with the virtual hand movements or with their unseen real hand movements. Due to the constant delay and resulting position mismatch between the virtual and real hands, only one of the hands’ (virtual or real) movements could be phase-matched to the target rhythm at any time point – the respective other hand was consequently incongruent with the target’s oscillatory phase, and effectively constituted a cross-modal distractor. This led to two different conditions and attentional task sets: “real hand”, which had participants ignore the visual information of the delayed virtual hand and focus only on proprioceptive information; and “virtual hand”, which had them ignore proprioception and focus only on visual information. Note that this task was pre-trained; i.e., we were not interested in learning effects but in sustained attention under cross-modal conflict (Limanowski & Friston, 2020a,b). The task instructions (‘VIRTUAL’ / ‘REAL’) were presented on screen before each block; furthermore, the colour of the fixation dot (pink or cyan, counterbalanced across participants) indicated the current instructed modality during movement.

Participants were instructed to maintain fixation on a centrally presented dot at all times. We recorded gaze data throughout each session with an eye tracker (Gazepoint GP3HD V2, https://www.gazept.com) at 60 Hz, positioned on the laptop at about 60 cm from the participants eyes.

Participants completed a brief training, and then 3 experimental sessions on separate days, separated by no less than 1 and no more than 2 weeks (Fig. 1D). Within each session, participants performed 8 trials (4 per VH and RH condition) of 65 target/grasping cycles (130 s), interspersed with 10 s rest period during which the instruction for the following trial was presented. The total task duration in each session was about 18.5 min.

### tACS

During each session, we either applied low-intensity sinusoidal tACS of 2 mA (peak-to-peak) at 10 Hz (“alpha”, cf. e.g. Clayton et al., 2019) or 20 Hz (“beta”, cf. e.g. Riosk et al., 2016) for 20 min or a sham protocol, which started with 30 seconds of current and then stopped. The order of stimulation sessions was counterbalanced, the experimenter was blind with respect to stimulation conditions while conducting the experiment and data analysis. The current was administered via conductive rubber electrodes placed into saline-soaked (NaCl 0.9%) sponges using a battery-operated stimulator (Starstim8, Neuroelectrics, Barcelona, Spain, www.neuroelectrics.com) controlled by the NIC2 software (Version 2.0, https://www.neuroelectrics.com/solutions/starstim/8). The electrodes were positioned over the occipital cortex (Oz) and centrally (Cz), using neoprene EEG caps fitted to the participant’s head circumference (Fig. 1C). Impedances were kept below 15 kΩ.

At the beginning of each session, participants filled out a self-assessment questionnaire. All participants stated ability to participate in the experiment, and no cognitive limitations. None of the participants reported to have consumed alcohol, nicotine, or any other drugs on the days of the experiment; except caffeine (up to 2 cups) and painkillers (only one participant’s session). After each experimental session, participants were asked to fill out a questionnaire about possible adverse effects of stimulation. In 8 out of 72 sessions the participants reported strong adverse effects of the simulation (5x “fatigue”, 1x “itching”, 1x “pain”, 1x “other”). In 3 sessions participants reported a strong effect of the stimulation on their general condition. In all other sessions no, or only mild to moderate sensations were reported. These values are in the range expected from previous reports and, therefore, can be interpreted as indicating successful participant blinding to stimulation condition (cf. Antal et al., 2017, 2025).

### Data pre-processing and analysis

The recorded hand movement trajectories (RH and VH) were normalized and epoched according to stimulation type (alpha, beta, sham) and task condition (RH vs VH). Task performance was quantified as the absolute phase shift between the target’s oscillatory cycle and the corresponding instructed hand trajectory (RH or VH, depending on task condition) using a Hilbert transform. Mean performance was obtained by averaging absolute phase shifts across grasping movements within each trial (excluding the first two auditory cycles as it typically takes participants one or two movements to engage with the task, cf. Limanowski & Friston, 2020a,b), and then averaging across the four trials of the same condition. To obtain performance variability, the standard deviation of the absolute phase shift was computed across samples within each trial and then averaged across trials of the same condition.

Eye-gaze position was recorded at 60 Hz as normalized x–y screen coordinates ([0, 1] range, with screen center at [0.5, 0.5]) along with a binary validity flag. Only valid gaze samples were retained. To evaluate the quality of fixation, we computed the Euclidean distance between each gaze sample and the screen center. Gaze data were epoched in the same manner as the hand-movement data to yield conditional estimates of average fixation and its variability.

We analyzed four behavioral outcomes (mean performance, performance variability, mean gaze-to-center distance, and gaze variability) using linear mixed-effects models (LMMs) implemented in Python (statsmodels, MixedLM). The dependent variable (DV) was modeled as a function of the experimental factors Factor A (stimulation type: alpha vs. beta vs. sham), Factor B (task: VH- vs. RH-matching), and Factor C (session order: 1–6). Factor C was included to evaluate whether the order of stimulation across sessions influenced the effects of the main factors of interest (A and B). All models included a participant-specific random intercept to account for within-subject dependence. Models were estimated by maximum likelihood (ML) to enable likelihood-ratio testing of nested fixed-effects structures. We fit a hierarchy of nested models to assess main effects and interactions (m0(base): DV ∼ FactorA * FactorB; m1: DV ∼ FactorA * FactorB + FactorC; m2: DV ∼ FactorA * FactorB + FactorA * FactorC + FactorB * FactorC; m3 (full): DV ∼ FactorA * FactorB * FactorC), where all factors were treated as categorical. Treatment coding was applied with sham stimulation as the reference level for Factor A, VH as the reference level for Factor B, and Session 1 as the reference level for Factor C. Models were compared using likelihood-ratio tests (LRTs), progressing from simpler to more complex fixed-effects structures (m0 → m1 → m2 → m3).

For each comparison, the likelihood-ratio statistic was computed as twice the difference in log-likelihoods and evaluated against a chi-square distribution with degrees of freedom equal to the number of additional fixed-effect parameters. This approach tested whether including interactions involving session order (Factor C) significantly improved model fit. After applying the model comparison hierarchically for the all four behavoural outcomes, we found that the addition of session order (Factor C) to the base model (m1 vs. m0) did not improve model fits (all ps > 0.58). Further models including interactions with Factor C (m2 vs. m1; m3 vs. m2) also showed no improvement (all ps > 0.07). Thus, the simpler model m0 was selected. For the selected model, we report fixed-effect coefficients (β), standard errors (SE), Wald z-statistics, two-sided p-values, and 95% confidence intervals. To follow up on significant effects obtained in the primary analysis, we calculated one-tailed paired t-tests.

## Results

Participants were, overall, able to align the instructed hand modality with the target in each condition, albeit lagging the target oscillation slightly (Fig. 2A). Hand-target matching performance was significantly better when participants had to align the VH than the RH (β = 0.32, SE = 0.09, p < .001, Fig. 2B), but there were no significant effects of alpha- or beta-tACS nor any significant interaction effects on average performance (all |βs| < 0.04, all ps > .68). There were no significant effects of alpha- or beta-tACS, instructed hand modality, or their interaction effects on performance variability (all |βs| < 0.05, all ps > .15, Fig. 2C).

**Figure 2:**
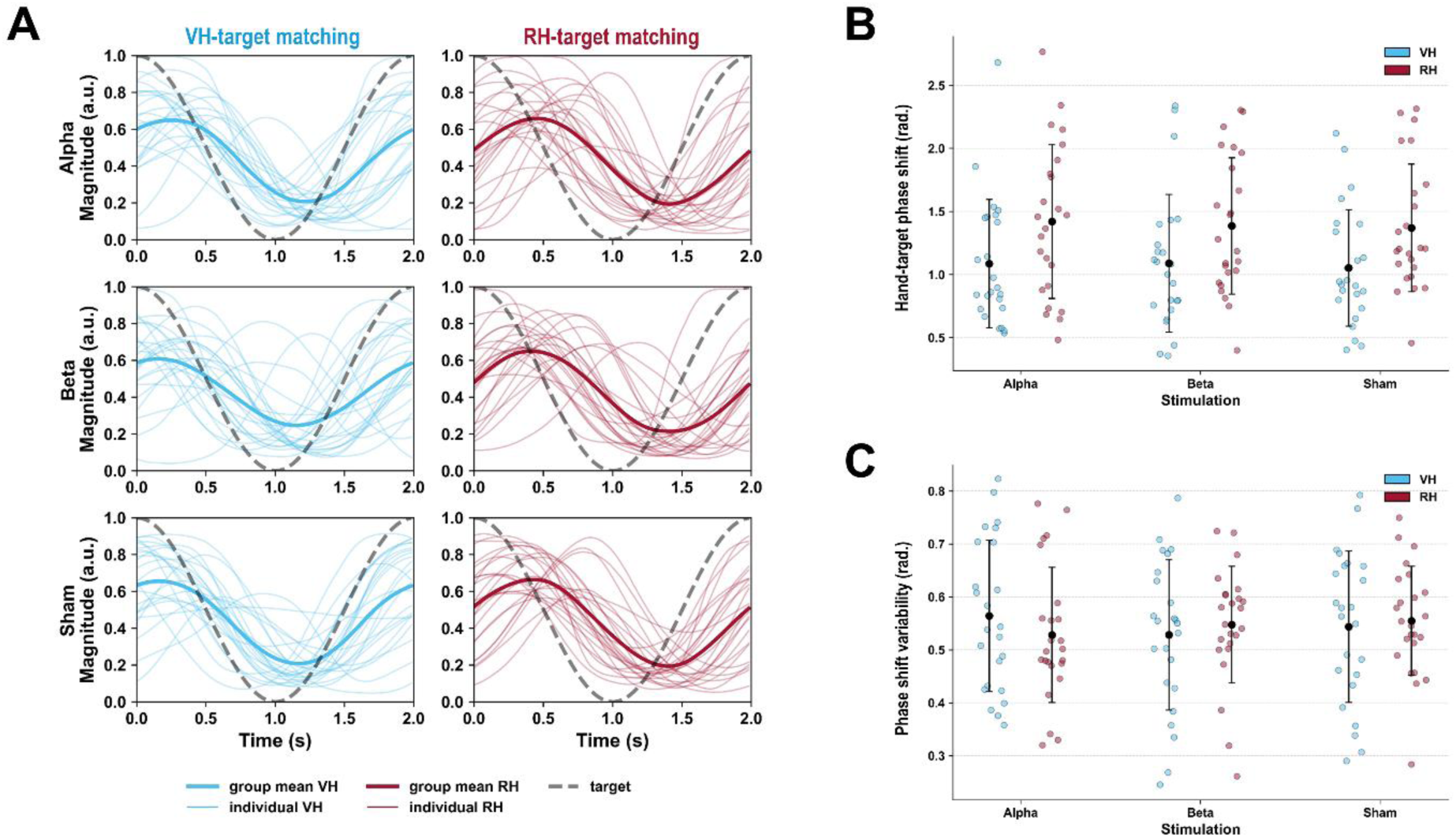
Hand-target matching performance. A: The colored curves show the grand-averaged grasping trajectories of the respective instructed hand modality (VH = virtual hand, blue lines; RH = real hand, red lines) relative to the target oscillation (grey dotted line). Thin lines are the individual participant averages. The non-instructed hand modality is not shown. **B-C:** Mean hand-target phase matching performance (B) and its average variability (C) per condition. Lower values indicate less phase shift between the instructed hand and the target oscillation and less variability, respectively; i.e., better performance in each case. The black dots and lines represent the means with associated standard deviations, the colored dots represent the individual participants’ averages.

**Table 1.** Results of the linear mixed-effects model for hand-target matching performance and variability (absolute phase shift, in radians). Fixed effects are reported with coefficient estimates (β), standard errors (SE), z values, p values, and 95% confidence intervals. Random effects are shown as variance components (n = 24).

| Predictor | $\beta$ | SE | z | p | 95% CI |
| --- | --- | --- | --- | --- | --- |
| *Hand-target matching performance* |  |  |  |  |  |
| Subject (intercept): Var = 0.171, SD = 0.188 |  |  |  |  |  |
| Intercept | 1.052 | 0.106 | 9.920 | <.001* | [0.844, 1.260] |
| Factor A (alpha vs. sham) | 0.034 | 0.091 | 0.370 | .711 | [-0.144, 0.212] |
| Factor A (beta vs. sham) | 0.037 | 0.091 | 0.412 | .681 | [-0.141, 0.216] |
| Factor B (rh vs. vh) | 0.317 | 0.091 | 3.489 | <.001* | [0.139, 0.495] |
| A (alpha) × B (rh) | 0.017 | 0.129 | 0.129 | .897 | [-0.235, 0.269] |
| A (beta) × B (rh) | -0.021 | 0.129 | -0.166 | .868 | [-0.273, 0.231] |
| *Hand-target matching variability* |  |  |  |  |  |
| Subject (intercept): Var = 0.010, SD = 0.043 |  |  |  |  |  |
| Intercept | 0.544 | 0.026 | 21.084 | <.001* | [0.493, 0.594] |
| Factor A (alpha vs. sham) | 0.020 | 0.023 | 0.885 | .376 | [-0.025, 0.065] |
| Factor A (beta vs. sham) | -0.015 | 0.023 | -0.677 | .499 | [-0.060, 0.029] |
| Factor B (rh vs. vh) | 0.011 | 0.023 | 0.481 | .631 | [-0.034, 0.056] |
| A (alpha) × B (rh) | -0.047 | 0.032 | -1.449 | .147 | [-0.110, 0.017] |
| A (beta) × B (rh) | 0.008 | 0.032 | 0.251 | .802 | [-0.055, 0.071] |

Gaze behavior was overall good; i.e., participants maintained fixation within minimal distance from the instructed fixation dot (Fig. 3A). There were no significant effects of alpha- or beta-tACS, instructed hand modality, nor their interaction on average fixation (all |βs| < 0.05, all ps > .12).

**Figure 3:**
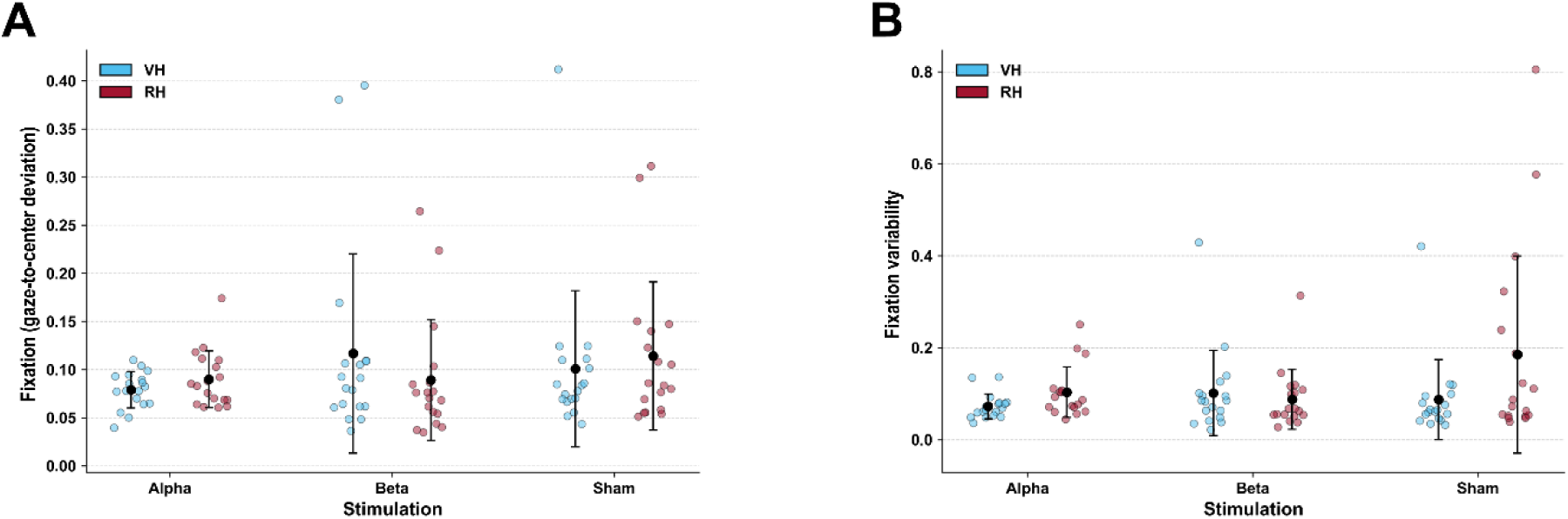
Gaze behavior. A: Average fixation performance per condition. Lower values indicate average gaze closer to the fixation dot. **B:** Average fixation variability per condition. Lower values indicate more consistent fixation throughout the respective condition. The black dots and lines represent the means with associated standard deviations, the colored dots represent the individual participants’ averages.

**Table 2.** Results of the linear mixed-effects model fixation performance and variability. Fixed effects are reported with coefficient estimates (β), standard errors (SE), z values, p values, and 95% confidence intervals. Random effects are shown as variance components (n = 18).

| Predictor | $\beta$ | SE | z | p | 95% CI |
| --- | --- | --- | --- | --- | --- |
| *Fixation performance* |  |  |  |  |  |
| Subject (intercept): Var = 0.001, SD = 0.013 |  |  |  |  |  |
| Intercept | 0.101 | 0.016 | 6.406 | <.001* | [0.070, 0.132] |
| Factor A (alpha vs. sham) | -0.022 | 0.018 | -1.188 | .235 | [-0.058, 0.014] |
| Factor A (beta vs. sham) | 0.016 | 0.018 | 0.853 | .394 | [-0.020, 0.052] |
| Factor B (rh vs. vh) | 0.013 | 0.018 | 0.715 | .474 | [-0.023, 0.049] |
| A (alpha) × B (rh) | -0.002 | 0.026 | -0.087 | .930 | [-0.054, 0.049] |
| A (beta) × B (rh) | -0.041 | 0.026 | -1.559 | .119 | [-0.092, 0.010] |
| *Fixation variability* |  |  |  |  |  |
| Subject (intercept): Var = 0.002, SD = 0.015 |  |  |  |  |  |
| Intercept | 0.087 | 0.025 | 3.513 | <.001* | [0.038, 0.136] |
| Factor A (alpha vs. sham) | -0.015 | 0.031 | -0.472 | .637 | [-0.076, 0.046] |
| Factor A (beta vs. sham) | 0.014 | 0.031 | 0.455 | .649 | [-0.047, 0.075] |
| Factor B (rh vs. vh) | 0.098 | 0.031 | 3.146 | .002* | [0.037, 0.159] |
| A (alpha) × B (rh) | -0.067 | 0.044 | -1.526 | .127 | [-0.154, 0.019] |
| A (beta) × B (rh) | -0.112 | 0.044 | -2.537 | .011* | [-0.198, -0.025] |

However, there was a significant effect of instructed hand modality on fixation variability (β = 0.10, SE = 0.03, p < .01); i.e., fixation was overall more variable in the RH than the VH condition (Fig. 3B). Furthermore, there was an interaction of instructed hand modality and beta-tACS (β = -0.11, SE = 0.04, p < .05), suggesting that the gaze variability effect above differed between the beta-tACS and sham conditions. Post-hoc t-tests confirmed that gaze variability during the RH condition was significantly lower under beta-tACS than sham stimulation (t(17) = 2.03, p < .05). There was no such effect of beta-tACS on gaze variability during the VH condition compared with sham (t(17) = -0.51, p = .69). Alpha-tACS reduced fixation variability relative to sham in the VH and RH conditions, but non-significantly.

## Discussion

Participants performed equally well across stimulation conditions, with similar performance (average and variability). Performance was overall better in the VH than the RH task (in line with our previous findings, Limanowski & Friston, 2020), which could indicate that monitoring the hand position for target alignment may be easier in the VH task, due to the relatively higher spatial acuity of vision than proprioception. However, this effect did not interact with stimulation type either. Thus, contrary to our expectations, neither alpha- nor beta-tACS significantly affected hand-target matching performance.

We observed an interaction effect on gaze behavior; i.e., fixation variability significantly decreased during beta-tACS relative to sham stimulation during the RH but not the VH condition (where variability was slightly higher under beta-tACS than sham). The average fixation performance was also slightly, but non-significantly better in RH task under beta-tACS than sham stimulation; again, in the VH task it was slightly higher under beta-tACS than sham. In comparison, alpha-tACS somewhat (non-significantly) improved average fixation and variability, but did not show the above interaction effect; i.e., it had the same (non-significant) effect on the VH and RH tasks compared with sham stimulation. In sum, beta-tACS seemed to selectively improve the participants’ ability to maintain central fixation in the RH condition, partly supporting our initial hypothesis.

Note that, in this kind of task, ignoring conflicting visual input is key to adequate RH task performance (Limanowski & Friston, 2020b), because incongruent visual movements are known to bias movement execution (Brass et al., 2001; Kilner et al., 2003; Heyes, 2011). Fixation variability was overall significantly higher in the RH than the VH task, which could indicate that the to-be-ignored, visual hand movements tended to capture attention and, therefore, produced unwanted “pursuit” eye movements. Conversely, more stable fixation could indicate that participants could better ignore these (visual) hand movements and, potentially, maintain sustained attention better onto proprioceptive movement feedback. That occipital beta-tACS facilitated maintaining central fixation in the RH condition could, therefore, support our assumption that augmenting beta oscillations over visual cortices may inhibit visual attention.

The fact that we only observed such effects under beta-, not alpha-tACS is somewhat surprising, given that visual attention and the control of (saccadic) eye movements has been linked to alpha oscillatory activity (Staudigl et al., 2017; Popov et al., 2017; Liu et al., 2023). Beta oscillations have mainly been linked to (oculo)motor control when recorded over motor cortices (e.g., Babapoor-Farrokhran et al., 2017; cf. Murthy & Fetz, 1992; Sanes & Donoghue, 1993). However, visual alpha and beta suppression has also been observed during smooth pursuit (Dunkley et al., 2015; cf. Churchland & Lisberger, 2005), where beta suppression might also have reflected processes related to motion perception per se (Dunkley et al., 2015). Furthermore, posterior beta suppression during visually guided action has been shown to co-vary with rhythmic movements, with the strongest suppression observed when attention to visual movement feedback was task-relevant (Wang & Limanowski, 2023). In line with these prior works, our result suggest that augmenting occipital beta oscillations via tACS facilitated ignoring visual movement feedback in conditions where it was a task-irrelevant distractor (RH task). This interpretation tentatively supports the general hypothesis that beta communicates top-down (attentional) control signals (Miller & Buschmann, 2013; Engel & Fries, 2010)

Our results should be interpreted in light of the following limitations. Firstly, there are several potential reasons for why we did not observe an effect of stimulation on manual performance: As we did not record brain activity, we cannot exclude the possibility that tACS simply did not modulate cortical oscillations. This question should be clarified by future work using concurrent electroencephalography recordings (cf. Ehrhardt et al., 2025). Furthermore, we stimulated at fixed frequencies. Brain stimulation generally may benefit from personalized stimulation parameters (cf. de Graaf et al., 2020; Fromm et al., 2025; Bergmann et al., 2025). Thus, future work should explore whether stimulating at participant-specific (e.g., peak alpha/beta) frequencies can yield the hypothesized behavioral effects. The eye tracking results were obtained using only 18 of 24 participants and need to be replicated in a larger sample. Future work should use more specialized study designs to test if the observed reduction in fixation variability is indeed related to visual suppression, e.g., by including psychophysical measures. This could open up an interesting avenue to explore potential benefits of beta-tACS on the attentional suppression of (visual) distractors during visuo-proprioceptive conflicts, for instance during action in virtual or remote environments.

In sum, our results suggest no direct effect of occipital alpha/beta-tACS on manual performance during visuomotor conflicts; but suggest that beta-tACS may improve oculomotor behavior related to sustained attention in the presence of to-be-ignored, visual distractors.

## Acknowledgments

We thank Juliane Füchtemann, Anna E. Fromm, Fanni Peters, Melanie Hertwig, and Zhenyu Wang for help with setup and data acquisition, and Jan Crucius for providing the eye tracker.

## Funding

Funding was provided by the Deutsche Forschungsgemeinschaft (DFG, German Research Foundation) – 426477764 to DA (AN 1103/3 − 1) and AF (FL 379/24 − 1), Research Unit 5429/1 (467143400) to AF (FL 379/34 − 1, FL 379/35 − 1) and DA (AN 1103/5 − 1), SFB1315/B03 (327654276) to AF, and 539593253 (to DA: AN 1103/6 − 1). The funders had no role in study design, data collection and analysis, decision to publish or preparation of the manuscript.

## Conflict of interest

None declared.

## Notes

### Competing Interest Statement

The authors have declared no competing interest.

## References

Antal, A., Alekseichuk, I., Bikson, M., Brockmöller, J., Brunoni, A. R., Chen, R., … & Paulus, W. (2017). Low intensity transcranial electric stimulation: safety, ethical, legal regulatory and application guidelines. Clinical neurophysiology, 128(9), 1774–1809.

Antal, A., Bjekić, J., Ganho-Ávila, A., Alekseichuk, I., Assecondi, S., Bergmann, T. O., … & Ziemann, U. (2025). Low intensity transcranial electric stimulation: Safety, ethical, legal regulatory and application guidelines (2017–2025: An update)–endorsed by the European Society for Brain Stimulation and by the International Federation for Clinical Neurophysiology. Clinical Neurophysiology, 2111436.

Arnal, L. H., Wyart, V., & Giraud, A.-L. (2011). Transitions in neural oscillations reflect prediction errors generated in audiovisual speech. Nature Neuroscience, 14(6), 797–801. 10.1038/nn.2810

Babapoor-Farrokhran, S., Vinck, M., Womelsdorf, T., & Everling, S. (2017). Theta and beta synchrony coordinate frontal eye fields and anterior cingulate cortex during sensorimotor mapping. Nature communications, 8(1), 13967.

Balslev, D., Christensen, L. O. D., Lee, J.-H., Law, I., Paulson, O. B., & Miall, R. C. (2004). Enhanced Accuracy in Novel Mirror Drawing after Repetitive Transcranial Magnetic Stimulation-Induced Proprioceptive Deafferentation. The Journal of Neuroscience, 24(43), 9698. 10.1523/JNEUROSCI.1738-04.2004

Bastos, A. M., Vezoli, J., Bosman, C. A., Schoffelen, J.-M., Oostenveld, R., Dowdall, J. R., De Weerd, P., Kennedy, H., & Fries, P. (2015). Visual Areas Exert Feedforward and Feedback Influences through Distinct Frequency Channels. Neuron, 85(2), 390–401. 10.1016/j.neuron.2014.12.018

Battaglini, L., Ghiani, A., Casco, C., & Ronconi, L. (2020). Parietal tACS at beta frequency improves vision in a crowding regime. NeuroImage, 208, 116451. 10.1016/j.neuroimage.2019.116451

Bauer, M., Kennett, S., & Driver, J. (2012). Attentional selection of location and modality in vision and touch modulates low-frequency activity in associated sensory cortices. Journal of Neurophysiology, 107(9), 2342–2351. 10.1152/jn.00973.2011

Bauer, M., Oostenveld, R., Peeters, M., & Fries, P. (2006). Tactile Spatial Attention Enhances Gamma-Band Activity in Somatosensory Cortex and Reduces Low-Frequency Activity in Parieto-Occipital Areas. Journal of Neuroscience, 26(2), 490–501. 10.1523/JNEUROSCI.5228-04.2006

Bauer, M., Stenner, M.-P., Friston, K. J., & Dolan, R. J. (2014). Attentional Modulation of Alpha/Beta and Gamma Oscillations Reflect Functionally Distinct Processes. Journal of Neuroscience, 34(48), 16117–16125. 10.1523/JNEUROSCI.3474-13.2014

Bergmann, T. O., Zoefel, B., Herrmann, C. S., Violante, I. R., Grossmann, N., Hassan, U., … & Hanslmayr, S. (2025). The tACS challenge: Does 10 Hz tACS rhythmically modulate visual perception in humans?(Stage 1 registered report accepted). Nature Human Behaviour.

Bernier, P.-M., Burle, B., Vidal, F., Hasbroucq, T., & Blouin, J. (2009). Direct Evidence for Cortical Suppression of Somatosensory Afferents during Visuomotor Adaptation. Cerebral Cortex, 19(9), 2106–2113. 10.1093/cercor/bhn233

Brass, M., Zysset, S., & von Cramon, D. Y. (2001). The Inhibition of Imitative Response Tendencies. NeuroImage, 14(6), 1416–1423. 10.1006/nimg.2001.0944

Buschman, T. J., & Miller, E. K. (2007). Top-Down Versus Bottom-Up Control of Attention in the Prefrontal and Posterior Parietal Cortices. Science, 315(5820), 1860–1862. 10.1126/science.1138071

Churchland, A. K., & Lisberger, S. G. (2005). Relationship between extraretinal component of firing rate and eye speed in area MST of macaque monkeys. Journal of neurophysiology, 94(4), 2416–2426.

Clayton, M. S., Yeung, N., & Cohen Kadosh, R. (2019). Electrical stimulation of alpha oscillations stabilizes performance on visual attention tasks. Journal of Experimental Psychology: General, 148(2), 203–220. 10.1037/xge0000502

de Graaf, T. A., Thomson, A., Janssens, S. E. W., van Bree, S., ten Oever, S., & Sack, A. T. (2020). Does alpha phase modulate visual target detection? Three experiments with tACS-phase-based stimulus presentation. European Journal of Neuroscience, 51(11), 2299–2313. 10.1111/ejn.14677

Donner, T. H., & Siegel, M. (2011). A framework for local cortical oscillation patterns. Trends in Cognitive Sciences, 15(5), 191–199. 10.1016/j.tics.2011.03.007

Dunkley, B. T., Freeman, T. C., Muthukumaraswamy, S. D., & Singh, K. D. (2015). Evidence that smooth pursuit velocity, not eye position, modulates alpha and beta oscillations in human middle temporal cortex. Human Brain Mapping, 36(12), 5220–5232.

Engel, A. K., & Fries, P. (2010). Beta-band oscillations—Signalling the status quo? Current Opinion in Neurobiology, 20(2), 156–165. 10.1016/j.conb.2010.02.015

Foxe, J. J., & Simpson, G. V. (2005). Biasing the brain’s attentional set: II. Effects of selective intersensory attentional deployments on subsequent sensory processing. Experimental Brain Research, 166(3), 393–401. 10.1007/s00221-005-2379-6

Foxe, J. J., Simpson, G. V., & Ahlfors, S. P. (1998). Parieto-occipital ∼1 0Hz activity reflects anticipatory state of visual attention mechanisms. NeuroReport, 9(17), 3929.

Fries, P., Reynolds, J. H., Rorie, A. E., & Desimone, R. (2001). Modulation of Oscillatory Neuronal Synchronization by Selective Visual Attention. Science, 291(5508), 1560–1563. 10.1126/science.1055465

Fromm, A. E., Trujillo-Llano, C., Grittner, U., Meinzer, M., Flöel, A., & Antonenko, D. (2025). Increased variability in response to transcranial direct current stimulation in healthy older compared to young adults: A systematic review and meta-analysis. Brain stimulation, 18(4), 1257–1265.

Fu, K.-M. G., Foxe, J. J., Murray, M. M., Higgins, B. A., Javitt, D. C., & Schroeder, C. E. (2001). Attention-dependent suppression of distracter visual input can be cross-modally cued as indexed by anticipatory parieto–occipital alpha-band oscillations. Cognitive Brain Research, 12(1), 145–152. 10.1016/S0926-6410(01)00034-9

Gundlach, C., Forschack, N., & Müller, M. M. (2024). Alpha-band fluctuations represent behaviorally relevant excitability changes as a consequence of top–down guided spatial attention in a probabilistic spatial cueing design. Imaging Neuroscience, 2, 1–24.

Haegens, S., Luther, L., & Jensen, O. (2012). Somatosensory Anticipatory Alpha Activity Increases to Suppress Distracting Input. Journal of Cognitive Neuroscience, 24(3), 677–685. 10.1162/jocn_a_00164

He, Q., Yang, X.-Y., Gong, B., Bi, K., & Fang, F. (2022). Boosting visual perceptual learning by transcranial alternating current stimulation over the visual cortex at alpha frequency. Brain Stimulation, 15(3), 546–553. 10.1016/j.brs.2022.02.018

Helfrich, R. F., Schneider, T. R., Rach, S., Trautmann-Lengsfeld, S. A., Engel, A. K., & Herrmann, C. S. (2014). Entrainment of Brain Oscillations by Transcranial Alternating Current Stimulation. Current Biology, 24(3), 333–339. 10.1016/j.cub.2013.12.041

Heuer, H., & Rapp, K. (2012). Adaptation to novel visuo-motor transformations: Further evidence of functional haptic neglect. Experimental Brain Research, 218(1), 129–140. 10.1007/s00221-012-3013-z

Heyes, C. (2011). Automatic imitation. Psychological Bulletin, 137(3), 463–483. 10.1037/a0022288

Hilla, Y., Link, F., & Sauseng, P. (2023). Alpha-tACS alters attentional control but not cognitive functions as video games do: A psychophysical investigation based on the theory of visual attention. European Journal of Neuroscience, 57(10), 1705–1722. 10.1111/ejn.15968

Kanai, R., Chaieb, L., Antal, A., Walsh, V., & Paulus, W. (2008). Frequency-Dependent Electrical Stimulation of the Visual Cortex. Current Biology, 18(23), 1839–1843. 10.1016/j.cub.2008.10.027

Kasten, F. H., Wendeln, T., Stecher, H. I., & Herrmann, C. S. (2020). Hemisphere-specific, differential effects of lateralized, occipital–parietal α- versus γ-tACS on endogenous but not exogenous visual-spatial attention. Scientific Reports, 10(1), 12270. 10.1038/s41598-020-68992-2

Kelly, S. P., Lalor, E. C., Reilly, R. B., & Foxe, J. J. (2006). Increases in Alpha Oscillatory Power Reflect an Active Retinotopic Mechanism for Distracter Suppression During Sustained Visuospatial Attention. Journal of Neurophysiology, 95(6), 3844–3851. 10.1152/jn.01234.2005

Kilner, J. M., Paulignan, Y., & Blakemore, S. J. (2003). An Interference Effect of Observed Biological Movement on Action. Current Biology, 13(6), 522–525. 10.1016/S0960-9822(03)00165-9

Lafleur, L.-P., Murray, A., Desforges, M., Pacheco-Barrios, K., Fregni, F., Tremblay, S., Saint-Amour, D., Lepage, J.-F., & Théoret, H. (2021). No aftereffects of high current density 10 Hz and 20 Hz tACS on sensorimotor alpha and beta oscillations. Scientific Reports, 11(1), 21416. 10.1038/s41598-021-00850-1

Lajoie, Y., Paillard, J., Teasdale, N., Bard, C., Fleury, M., Forget, R., & Lamarre, Y. (1992). Mirror drawing in a deafferented patient and normal subjects. Neurology, 42(5), 1104–1104. 10.1212/WNL.42.5.1104

Lebar, N., Danna, J., Moré, S., Mouchnino, L., & Blouin, J. (2017). On the neural basis of sensory weighting: Alpha, beta and gamma modulations during complex movements. NeuroImage, 150, 200–212. 10.1016/j.neuroimage.2017.02.043

Lee, J. H., Whittington, M. A., & Kopell, N. J. (2013). Top-Down Beta Rhythms Support Selective Attention via Interlaminar Interaction: A Model. PLOS Computational Biology, 9(8), e1003164. 10.1371/journal.pcbi.1003164

Liu, B., Nobre, A. C., & van Ede, F. (2023). Microsaccades transiently lateralise EEG alpha activity. Progress in Neurobiology, 224, 102433.

Limanowski, J. (2022). Precision control for a flexible body representation. Neuroscience & Biobehavioral Reviews, 134, 104401. 10.1016/j.neubiorev.2021.10.023

Limanowski, J., & Friston, K. (2020a). Active inference under visuo-proprioceptive conflict: Simulation and empirical results. Scientific Reports, 10(1), 4010. 10.1038/s41598-020-61097-w

Limanowski, J., & Friston, K. (2020b). Attentional Modulation of Vision Versus Proprioception During Action. Cerebral Cortex, 30(3), 1637–1648. 10.1093/cercor/bhz192

Limanowski, J., Litvak, V., & Friston, K. (2020). Cortical beta oscillations reflect the contextual gating of visual action feedback. NeuroImage, 222, 117267. 10.1016/j.neuroimage.2020.117267

Miller, E. K., & Buschman, T. J. (2013). Cortical circuits for the control of attention. Current opinion in neurobiology, 23(2), 216–222.

Murthy, V. N., & Fetz, E. E. (1992). Coherent 25-to 35-Hz oscillations in the sensorimotor cortex of awake behaving monkeys. Proceedings of the National Academy of Sciences, 89(12), 5670–5674.

Nakazono, H., Ogata, K., Kuroda, T., & Tobimatsu, S. (2016). Phase and Frequency-Dependent Effects of Transcranial Alternating Current Stimulation on Motor Cortical Excitability. PLOS ONE, 11(9), e0162521. 10.1371/journal.pone.0162521

Nakazono, H., Ogata, K., Takeda, A., Yamada, E., Kimura, T., & Tobimatsu, S. (2020). Transcranial alternating current stimulation of α but not β frequency sharpens multiple visual functions. Brain Stimulation, 13(2), 343–352. 10.1016/j.brs.2019.10.022

Pogosyan, A., Gaynor, L. D., Eusebio, A., & Brown, P. (2009). Boosting Cortical Activity at Beta-Band Frequencies Slows Movement in Humans. Current Biology, 19(19), 1637–1641. 10.1016/j.cub.2009.07.074

Popov, T., Kastner, S., & Jensen, O. (2017). FEF-controlled alpha delay activity precedes stimulus-induced gamma-band activity in visual cortex. Journal of Neuroscience, 37(15), 4117–4127.

Rjosk, V., Kaminski, E., Hoff, M., Gundlach, C., Villringer, A., Sehm, B., & Ragert, P. (2016). Transcranial Alternating Current Stimulation at Beta Frequency: Lack of Immediate Effects on Excitation and Interhemispheric Inhibition of the Human Motor Cortex. Frontiers in Human Neuroscience, 10. 10.3389/fnhum.2016.00560

Roshchupkina, L., Stee, W., & Peigneux, P. (2020). Beta-tACS does not impact the dynamics of motor memory consolidation. *Brain Stimulation: Basic*, Translational, and Clinical Research in Neuromodulation, 13(6), 1489–1490. 10.1016/j.brs.2020.08.012

Sanes, J. N., & Donoghue, J. P. (1993). Oscillations in local field potentials of the primate motor cortex during voluntary movement. Proceedings of the National Academy of Sciences, 90(10), 4470–4474.

Schroeder, C. E., & Lakatos, P. (2009). Low-frequency neuronal oscillations as instruments of sensory selection. Trends in Neurosciences, 32(1), 9–18. 10.1016/j.tins.2008.09.012

Schuhmann, T., Kemmerer, S. K., Duecker, F., Graaf, T. A. de, Oever, S. ten, Weerd, P. D., & Sack, A. T. (2019). Left parietal tACS at alpha frequency induces a shift of visuospatial attention. PLOS ONE, 14(11), e0217729. 10.1371/journal.pone.0217729

Schutter, D. J. L. G., & Hortensius, R. (2011). Brain oscillations and frequency-dependent modulation of cortical excitability. Brain Stimulation, 4(2), 97–103. 10.1016/j.brs.2010.07.002

Spitzer, B., & Haegens, S. (2017). Beyond the Status Quo: A Role for Beta Oscillations in Endogenous Content (Re)Activation. eNeuro, 4(4). 10.1523/ENEURO.0170-17.2017

Staudigl, T., Hartl, E., Noachtar, S., Doeller, C. F., & Jensen, O. (2017). Saccades are phase-locked to alpha oscillations in the occipital and medial temporal lobe during successful memory encoding. PLoS biology, 15(12), e2003404.

Wang, P., & Limanowski, J. (2023). Phasic modulation of beta power at movement-related frequencies during visuomotor conflict. Journal of Neurophysiology, 130(5), 1367–1372. 10.1152/jn.00338.2023

Zaehle, T., Rach, S., & Herrmann, C. S. (2010). Transcranial Alternating Current Stimulation Enhances Individual Alpha Activity in Human EEG. PLOS ONE, 5(11), e13766. 10.1371/journal.pone.0013766

